# Single-cell transcriptomics reveals immune suppression and cell states predictive of patient outcomes in rhabdomyosarcoma

**DOI:** 10.1101/2022.07.15.497944

**Authors:** Jeff DeMartino, Michael T. Meister, Lindy Visser, Mariël Brok, Marian J. A. Groot Koerkamp, Laura S. Hiemcke-Jiwa, Terezinha de Souza, Johannes H. M. Merks, Frank C. P. Holstege, Thanasis Margaritis, Jarno Drost

## Abstract

Paediatric rhabdomyosarcoma (RMS) is a soft tissue malignancy of mesenchymal origin which is thought to arise as a consequence of derailed myogenic differentiation. Despite intensive treatment regimens, the prognosis for high-risk patients remains dismal. The cellular differentiation states underlying RMS and how these relate to patient outcomes remain largely elusive. Here, we used single-cell mRNA-sequencing to generate a transcriptomic atlas of RMS. Analysis of the RMS tumour niche revealed evidence of an immunosuppressive microenvironment. We also identified an interaction between NECTIN3 and TIGIT, specific to the more aggressive fusion-positive (FP) RMS subtype, as a putative cause of tumour-induced T-cell dysfunction. In malignant RMS cells we defined transcriptional programs reflective of normal myogenic differentiation. Furthermore, we showed that these cellular differentiation states are predictive of patient outcomes in both FP RMS and the more clinically homogenous fusion-negative subtype. Our study reveals the potential of therapies targeting the immune microenvironment of RMS and suggests that assessing tumour differentiation states may enable a more refined risk stratification.

## Main

Rhabdomyosarcoma (RMS) is the most commonly diagnosed soft tissue sarcoma (STS) in children and adolescents, accounting for approximately 3.5% of all paediatric malignancies^1^. Several characteristics, including expression of the myogenic regulatory transcription factors *MYOD1* and *MYOG*^2^ and the presence of rhabdomyoblasts^3^ (cells reminiscent of terminally differentiating myocytes) point to RMS being the result of impaired skeletal muscle myogenesis. However, the disease may also arise at body sites devoid of skeletal muscle, and RMS models of non-myogenic origin have been described^4^. Despite intense, multimodal treatment strategies, outcomes remain dismal for patients with high-risk or metastatic disease, the latter of which exhibits a long-term overall survival rate (OS) of approximately 30%^5^. This emphasizes the need to improve our understanding of RMS tumour biology to enable the development of novel therapeutic approaches.

Historically, RMS has been divided into two main subtypes, alveolar and embryonal, based on histological features of tumours^6^. However, recent work has shown that the molecular classification as either fusion positive (FP) or fusion negative (FN) is a more powerful prognostic indicator^7,8^. FP RMS is characterized by recurrent chromosomal translocations resulting in the expression of a chimeric fusion protein containing the DNA binding domains of either PAX3 or PAX7, both key transcriptional regulators of normal myogenesis^9^, coupled to a strong transactivation domain, most often that of FOXO1^10,11^. The genetic lesions driving FN RMS, on the other hand, are diverse and may include mutations in signal transduction pathways (especially RAS and PI3K), cell cycle regulators and the P53 pathway, among others^12^. Notably, FP RMS carries a significantly worse prognosis than FN RMS, and is more often metastatic at diagnosis^8^.

Beyond the inter-tumoral genetic heterogeneity characteristic of RMS, it has been recognized that there exists a degree of heterogeneity within tumours, as exemplified by the diversity in cellular morphology^13^ and variation in immunohistochemical staining for myogenic markers^14^. However, the characteristics and clinical implications of this heterogeneity remain unclear. Additionally, the composition of the tumour microenvironment (TME) and the interplay between malignant cells and the TME have not been comprehensively profiled. Here, we compile a single-cell transcriptomic atlas comprising both FN and FP RMS and find distinct differences in cellular composition and single-cell differentiation states between and within subtypes that relate to clinical outcomes and suggest potential immunotherapeutic interventions.

## Results

### A single-cell atlas of paediatric RMS tumours

We implemented a protocol for performing plate-based single-cell mRNA-sequencing^15^ (SORT-seq) on viably frozen RMS tumour samples (Fig. 1a and Extended Data Fig. 1a). Opting for a plate-based method allowed for the generation of high-quality single-cell transcriptomes from samples with low viability (including pre-treated samples) or where limited material was available (e.g., small needle biopsies). From our cohort of 19 RMS samples, encompassing the major molecular and histological subtypes (FP, FN, alveolar and embryonal, Fig. 1b,c and Supplementary Table S1), we obtained 7,364 high quality single-cell transcriptomes (median of 420 per sample) which passed quality thresholds.

**Figure 1:**
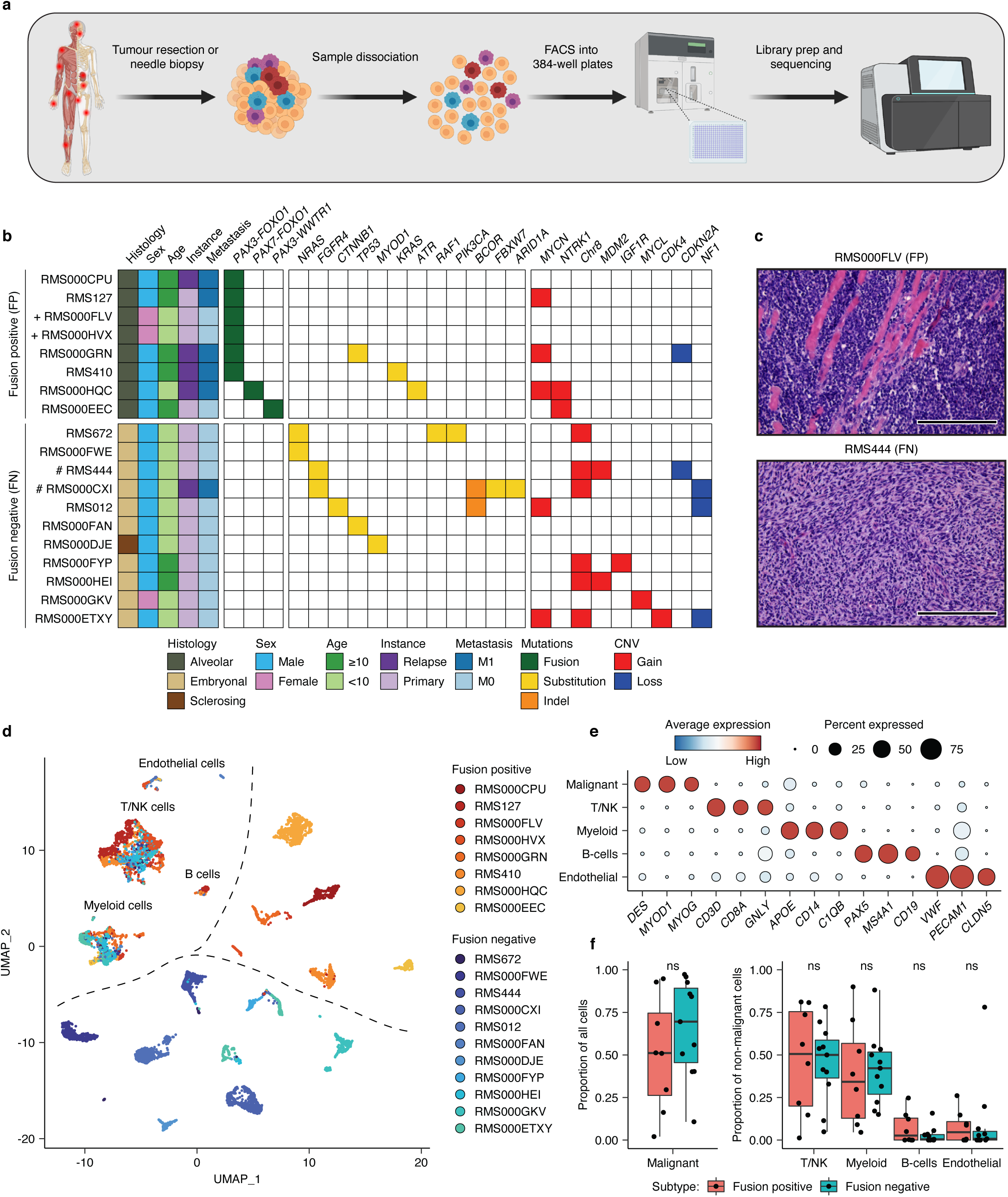
Single-cell transcriptomic atlas of RMS tumours. **a**, Schematic representation of the sample processing workflow used to generate scRNA-seq data. Created with BioRender. **b**, Overview of RMS sample cohort, including patient clinical characteristics, as well as a summary of relevant mutations and copy number variants (CNV) in tumours, defined using bulk DNA sequencing. (+) and (#) indicate independent samples derived from the same patient. **c**, Representative haematoxylin and eosin (H&E) stained tumour sections depicting the two major RMS histological subtypes (alveolar and embryonal) in this cohort. Scale bars are equivalent to 200*µ*m. **d**, UMAP projection of single-cell RMS transcriptomes (n = 7,364) coloured by sample. **e**, Dot plot depicting the average scaled gene expression of selected marker genes for each annotated cell type (dot colour). Dot size corresponds to the percentage of cells expressing each gene. **f**, Boxplots comparing the proportion of malignant cells (left panel) and of each non-malignant cell type (right panel) between molecular subtypes. ns = not significant (*p* > 0.05, Student’s t-test).

To distinguish RMS cells from non-malignant cell types comprising the TME, two complementary approaches were employed. First, the similarity between each single-cell transcriptome and a reference collection of bulk transcriptomes derived from healthy cell types and RMS tumours was assessed using *SingleR*^16^ (see Methods). Clustering of the resulting similarity scores revealed a clear distinction between cells with a high correspondence to bulk RMS tumours (malignant cells) and those which resembled one of several immune or stromal cell types (Extended Data Fig. 1b). Second, single-cell copy number variation (CNV) profiles were inferred and clustered on a per tumour basis. In all tumours, cells harbouring coherent whole and sub-chromosomal CNVs (malignant cells) could be distinguished from those which appear to be copy neutral (Extended Data Fig. 2). In general, single-cell derived CNV profiles were highly similar to those defined by DNA sequencing of bulk tumour samples (Extended Data Fig. 2). Cells classified as “malignant” or “normal” using both methods were retained, while divergently classified cells were excluded from further analysis. The median percentage of malignant cells per sample was 56%, though this varied widely (2%-97%), and did not differ significantly between molecular subtypes (Fig. 1f and Extended Data Fig. 1c). Putative malignant cells expressed high levels of classical RMS marker genes *DES, MYOD1* and *MYOG*, as expected (Fig. e). *SingleR* cell-type similarity scores and the expression of known marker genes were used to discern the identities of non-malignant cells (Fig. 1e and Extended Data Fig. 1b). As with the overall percentage of malignant cells, the proportion of each non-malignant cell type varied extensively between tumours but did not differ significantly based on fusion status (Fig. 1f and Extended Data Fig. 1c). Projecting the classified single-cell transcriptomes in Uniform Manifold Approximation and Projection (UMAP) space revealed that inter-tumoral heterogeneity and molecular subtype classification (FN or FP) drove the clustering of malignant cells, while non-malignant cells clustered by cell type (Fig. 1d), as has previously been described for other tumour entities^17–20^.

### Characterization of the RMS microenvironment reveals general and subtype-specific immune dysfunction

To explore the composition and functional characteristics of tumour-infiltrating immune cells, graph-based clustering was performed on the myeloid and T/NK compartments (Fig. 2b,d). Examination of marker gene expression in the myeloid clusters revealed the presence of undifferentiated (M0) and differentiated (Mq) macrophages, as well as conventional (cDC) and plasmacytoid (pDC) dendritic cells (Fig. 2b and Extended Data Fig. 3a). Scoring differentiated macrophages for M1/M2-specific gene signatures^21^ indicated that they existed predominantly in the M2 state (Fig. 2c), which has been associated with several pro-tumorigenic functions including the suppression of inflammation and promotion of angiogenesis^22^. Among the T/NK cell clusters, several subtypes could be discerned including naïve and gamma delta (ψ8) T cells, regulatory T cells (Tregs), cytotoxic (CD8+) T cells and multiple subtypes of CD4+ T helper cells (IL7R+ and ISG+) (Fig. 2d and Extended Data Fig. 3b). IHC for immune cell markers confirmed the presence of infiltrating T cells in RMS tissues (Extended Data Fig. 1d). Interestingly, interferon-stimulated T helper cells (ISG+) were found almost exclusively in FN tumours, which may reflect a higher degree of immunogenicity (Extended Data Fig. 3b).

**Figure 2:**
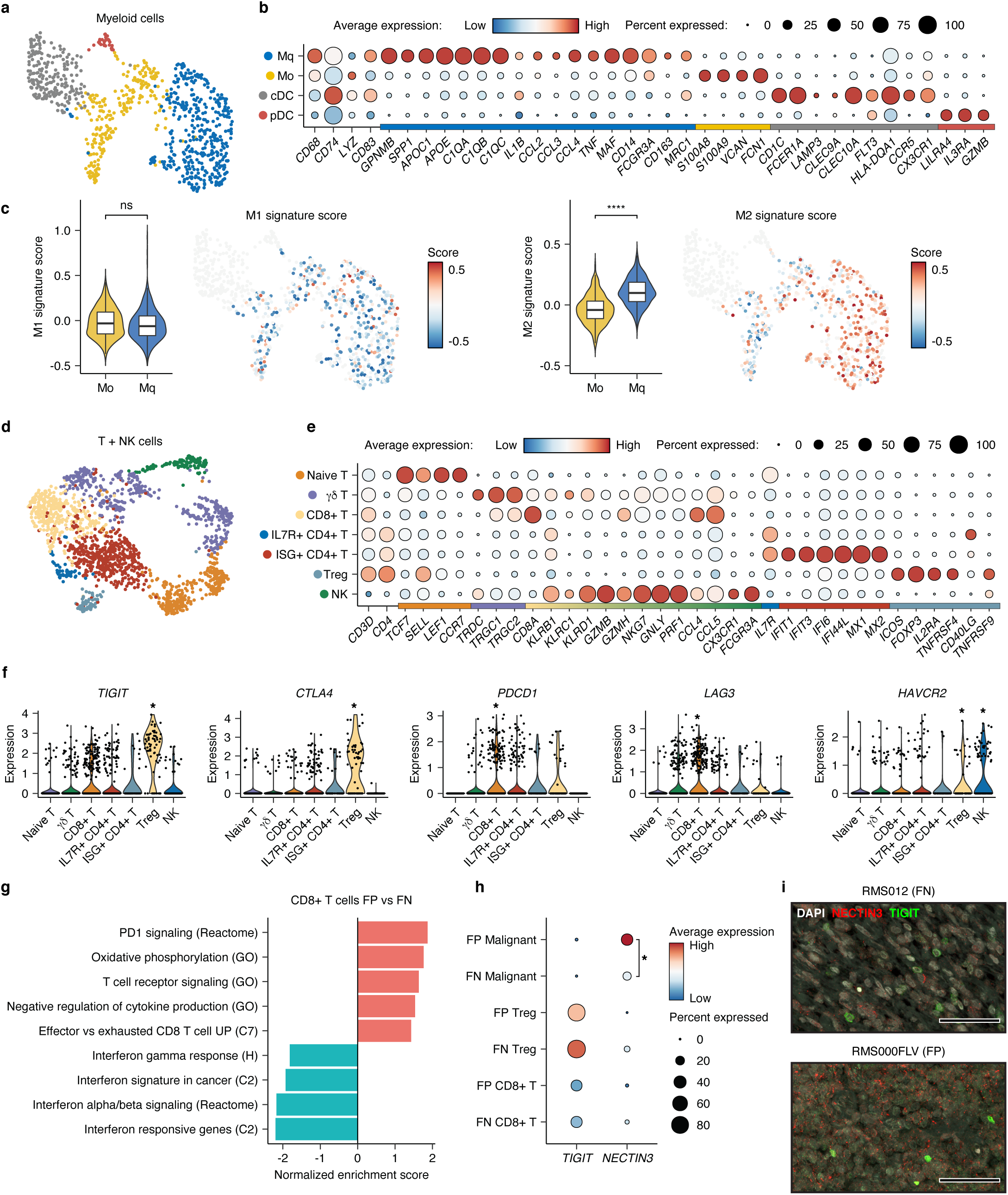
Characterization of the RMS immune microenvironment. **a**, UMAP projection of myeloid cells, coloured by cluster assignment. **b**, Dot plot depicting the average expression of selected cell type-specific genes. Dot size corresponds to the percentage of cells expressing each marker. Colour bar on the x-axis indicates for which cluster each gene is specific to. **c**, Violin and UMAP plots showing the distribution of M1 (left panel) and M2 (right panel) signature scores in undifferentiated (M0) and differentiated (MФ) macrophages. ns = not significant (p > 0.05, Student’s T-test), **** indicates p < 0.0001 (Student’s T-test). Non-macrophage cells are coloured grey on UMAP plots. **d**, UMAP projection of T and NK cells, coloured by cluster assignment. **e**, Dot plot depicting the average expression of selected cell type-specific genes. Dot size corresponds to the percentage of cells expressing each marker. Colour bar on the x-axis indicates the cluster specificity for each gene. **f**, Violin plots showing the expression of selected immune checkpoint molecules within T/NK subsets. * Indicates differential expression (Log2 FC > 0.25 and p < 0.05, Wilcoxon rank sum test). **g**, Normalized enrichment scores (NES) of selected gene sets, as determined by gene set enrichment analysis (GSEA) comparing CD8+ T cells between RMS subtypes. Codes in parenthesis indicate the database from which the gene set derives (H, C2 and C7 correspond to MSigDB collections). **h**, Dot plot depicting the average expression of the *TIGIT* and *NECTIN3* genes in selected cell types (per RMS subtype). Dot size corresponds to the percentage of cells expressing each marker. * Indicates differential expression (LogFC > 0.25 and p < 0.05, Wilcoxon rank sum test). **i**, Representative immunofluorescence (IF) microscopy images depicting the expression of TIGIT (green) and NECTIN3 (red), along with DAPI counterstaining (grey), in RMS tissue sections from FN and FP tumours. Scale bars equivalent to 50*µ*m.

Within T cell subgroups, the expression of several genes encoding molecules associated with immune dysfunction and the suppression of immune responses ^23^ was observed, including *LAG3* and *PDCD1* (PD1) in CD8+ T cells, *CTLA4* and *TIGIT* in Tregs and *HAVCR2* in Tregs and NK cells (Fig. 2f). Strikingly, gene set enrichment analysis (GSEA) comparing CD8+ T cells between RMS subtypes indicated that dysfunction was more prevalent in FP samples, which were enriched for gene sets related to PD-1 signalling, oxidative phosphorylation and T cell exhaustion, while cells from FN tumours were enriched for interferon response and stimulation signatures (Fig. 2g). To define putative cell-cell interactions regulating immune dysfunction, we used *CellChat*^24^ to model ligand-receptor interactions between malignant cells, per subtype, and cell types within the TME. This analysis highlighted a putative interaction specific to FP tumours between NECTIN3 expressed on malignant cells, and the TIGIT receptor on Tregs and CD8+ T cells (Extended Data Fig. 3c). The specificity of this interaction was due to the significantly higher expression of *NECTIN3* in FP tumour cells, while the expression of *TIGIT* in Tregs and CD8+ T cells was comparable between subtypes (Fig. 2h). Supporting this finding, immunofluorescence microscopy (IF) of tumour tissues revealed the presence of TIGIT-positive cells in both subtypes, while a more prevalent staining pattern of NECTIN3 was observed in FP RMS (Fig. 2i). Taken together, analysis of the TME in RMS highlighted evidence of general immune dysfunction, indicated by the prevalence of M2 macrophages, as well as a putative FP-specific T-cell exhaustion phenotype which may be partly regulated, by the interaction between NECTIN3 and TIGIT.

### Malignant cell states in RMS mirror normal myogenic differentiation

While it has been proposed that RMS tumours arise as a result of myogenic differentiation gone awry, the identification of the precise developmental origin(s) of RMS remains an active area of investigation^25^. To place RMS tumour cells within the context of normal myogenic differentiation, a series of logistic regression models were trained, as previously described^26^, to predict the similarity of malignant single-cell transcriptomes to the main cell types defined by a recently published single-cell atlas of human pre- and post-natal myogenesis^27^. This analysis showed that, on average, FN RMS cells resembled both myogenic progenitors and myogenic mesenchymal cells, while FP cells most closely corresponded to committed myoblasts (Extended Data Fig. 4a). This is in line with the notion that FN tumours often exhibit an undifferentiated “embryonal” histology, while FP more widely express the key myogenic regulatory factors *MYOD1* and *MYOG*^6^ (Fig. 1b,c and Extended Data Fig. 4b). However, when analysing at single-cell resolution we found that individual cells from each subtype and tumour spanned the spectrum of myogenic differentiation, indicating that there exists large-scale intra-, as well as inter-tumoral heterogeneity in cellular differentiation states (Extended Data Fig. 4a).

### NMF-defined differentiation trajectories in FN RMS reflect early myogenesis

To probe the prospective sources of heterogeneity, non-negative matrix factorization (NMF) was applied, independently per molecular subtype, to define the underlying transcriptional programs active in malignant cells from each of the tumours in our RMS scRNA-seq cohort (see Methods). In FN RMS samples, this analysis revealed three clusters of highly correlated transcriptional programs, which we merged into three meta-programs (Fig. 3a-left panel). Notably, the constituent programs underlying each meta-program were derived from several tumour samples, indicating that clustering was not driven by inter-tumoral heterogeneity. To interpret the biological relevance of each meta-program, we assessed the expression of their top weighted genes (Fig. 3a-right panel and Supplementary Table S2). The first program, which we termed “mesenchymal”, was enriched for genes related to extracellular matrix (ECM) organization, including *FN1, TGFBI* and several collagen-encoding genes, among others (Extended Data Fig. 5a). The second program, referred to as the “progenitor-like” program included genes expressed during early myogenesis^27^, such as *FGFR4* and *GPC3*, as well as markers of proliferation, including *MKI67* and *TOP2A*. Finally, the “myogenic” program, was characterized by genes involved in the regulation of myogenic differentiation, including *MYOD1, MYOG, MEF2C* and *CDH15* as well as genes encoding structural and functional components of terminally differentiated striated muscle, such as *TTN* and *CKM*. Scoring FN cells for each meta-program revealed that expression of the myogenic and mesenchymal programs was mutually exclusive, while expression of the highly proliferative progenitor-like state was restricted to cells which scored low for the mesenchymal as well as the myogenic programs (Fig. 3b). These patterns were corroborated using a dataset from a recently published independent single-nucleus RNA-seq cohort of RMS tumours^28^ (Extended Data Fig. 5b).

**Figure 3:**
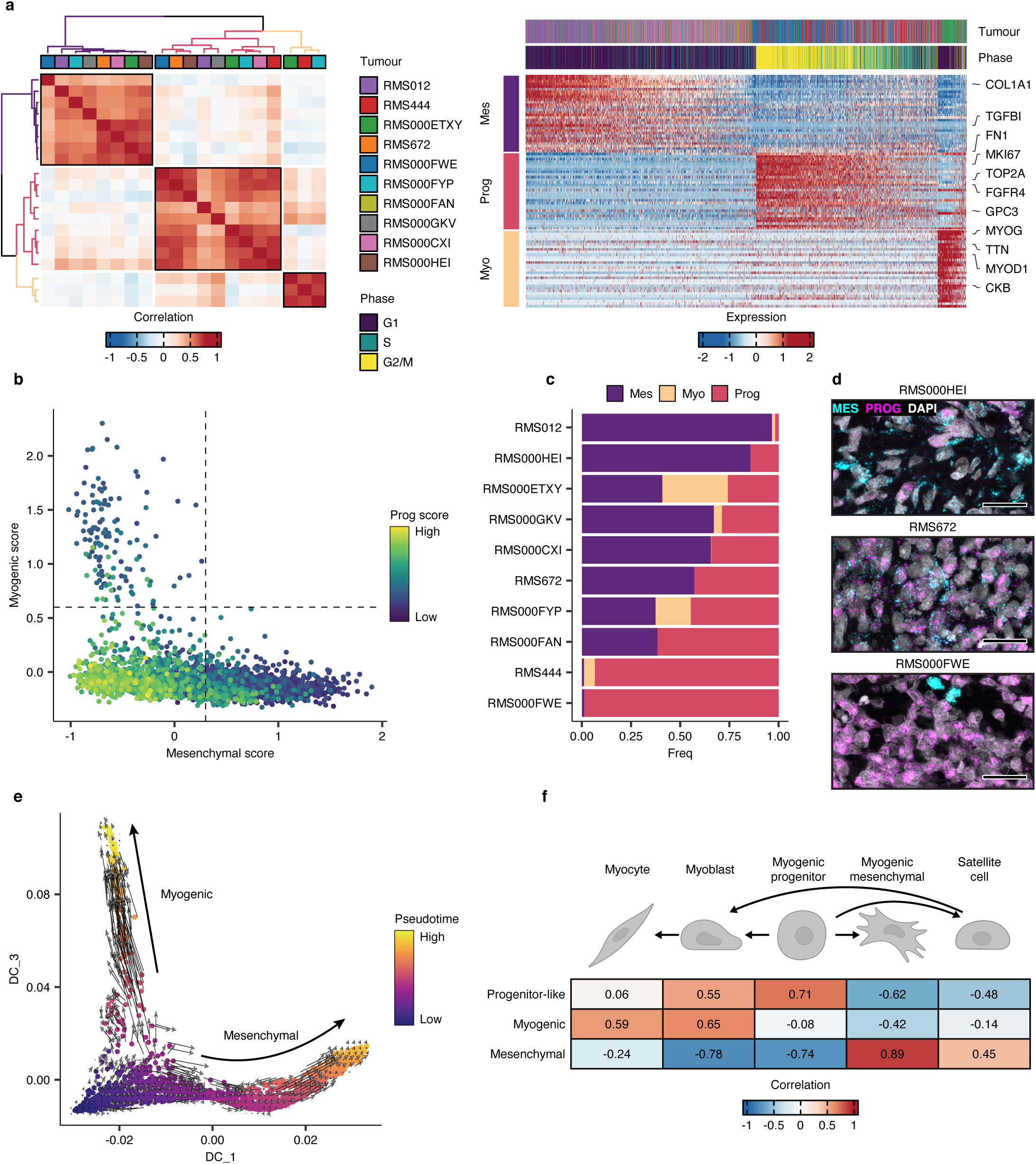
NMF defines malignant cell states in FN RMS tumours. **a**, Left panel: Heatmap showing the pairwise Pearson correlations between all NMF-defined transcriptional programs in FN samples. The tumour sample from which each transcriptional program derived is shown in the colour bar. Meta-program clusters are delineated by black boxes and colouring of the dendrograms. Right panel: Scaled expression of the top 30 genes per meta-program across all FN cells. The corresponding tumour sample and inferred cell cycle phase of each cell are displayed in the top annotation track. Representative genes from each meta-program are labelled. **b**, Scatterplot depicting the mesenchymal (x-axis), myogenic (y-axis) and proliferative (point colour) meta-program scores. Dotted lines correspond to the cut-offs used to define discrete cell states. **c**, Proportion of cells within each discrete state, per FN tumour. **d**, Representative RNA fluorescence *in-situ* hybridization (RNA-FISH) images depicting the expression of mesenchymal (MES = *TGFBI*) and progenitor-like (PROG = *FGFR4*) cell state marker genes in FN tissue samples. DAPI counterstaining shown in grey. Scale bars equivalent to 25*µ*m. **e**, Diffusion maps projection of FN RMS single cells, coloured by pseudotime value, overlaid with the RNA velocity vector field. **f**, Heatmap depicting the Pearson correlations between cell-state scores, and the logistic regression-defined similarity scores (logits) for each normal myogenic cell type. Myogenic differentiation schematic was created with BioRender.

Meta-program scores were then used to define the discrete “state” of each cell. This analysis revealed a high degree of variation between tumours in the distribution of cell states (Fig. 3c). Interestingly, some tumours (e.g., RMS012 and RMS000HEI) were dominated by mesenchymal-state cells, while others (e.g., RMS444 and RMS000FWE) almost exclusively contained progenitor-like-state cells. RNA fluorescence *in-situ* hybridization (RNA-FISH) was used to validate the presence of each cell-state and the distribution of the progenitor-like and mesenchymal states within individual tumours (Fig. 3d and Extended Data Fig. 5c). To investigate the hierarchy of cell states in FN RMS, the data was modelled as a differentiation trajectory by projecting single-cell transcriptomes in diffusion maps space and using pseudotime and RNA velocity to assess directionality (Fig. 3e and Methods). This analysis suggested that cells transition from the highly proliferative progenitor-like state into the more differentiated mesenchymal or myogenic states. Variation in differentiation status was also evident when comparing the malignant cell-state scores with the similarity scores to normal myogenic cell types. This showed that the progenitor-like score correlated strongly with undifferentiated myogenic progenitors, while the mesenchymal and myogenic scores with more differentiated cell types, namely myogenic mesenchymal cells, and myoblasts/myocytes, respectively (Fig. 3f). Together, these data show that transcriptional cell-states in FN RMS cells can be organized in a differentiation trajectory mirroring that of early myogenic differentiation, where progenitor-like cells can give rise to terminally differentiating myoblasts, or those progressing toward myogenic mesenchymal cells.

### Differentiation states in FP RMS mirror skeletal muscle regeneration

Extending the NMF analysis to FP RMS also revealed three meta-programs, as defined by correlating transcriptional programs across tumour samples (Fig. 4a-left panel). The proliferative program consisted almost entirely of genes involved in mitotic cell processes, including *MKI67, TOP2A* and *CENPE*, among others (Extended Data Fig. 6a). As expected, nearly all cells inferred to be in S or G2/M phases scored high for this meta-program (Fig. 4a-right panel and Supplementary Table S3). The myogenic program was marked by expression of terminal myogenic differentiation genes, such as *MYOG, TTN* and *CKB*. Finally, the program termed “satellite cell-like” (SC-like) was characterized by expression of the *NOTCH3* receptor gene, Notch pathway targets, including *HEY1* and *HES1*, and genes encoding type V and VI collagens. These genes are known to play roles in the context-specific regulation of quiescence, self-renewal, and activation in muscle-resident satellite cells^29,30^. Scoring single-cells for each meta-program revealed a mutually exclusive relationship between the myogenic and SC-like programs, while the proliferative program did not correlate with either and was, in general, restricted to cells scoring low for the two former programs (Fig. 4b). Again, the relationship between meta-program scores was confirmed in an independent dataset (Extended Data Fig 6b).

**Figure 4:**
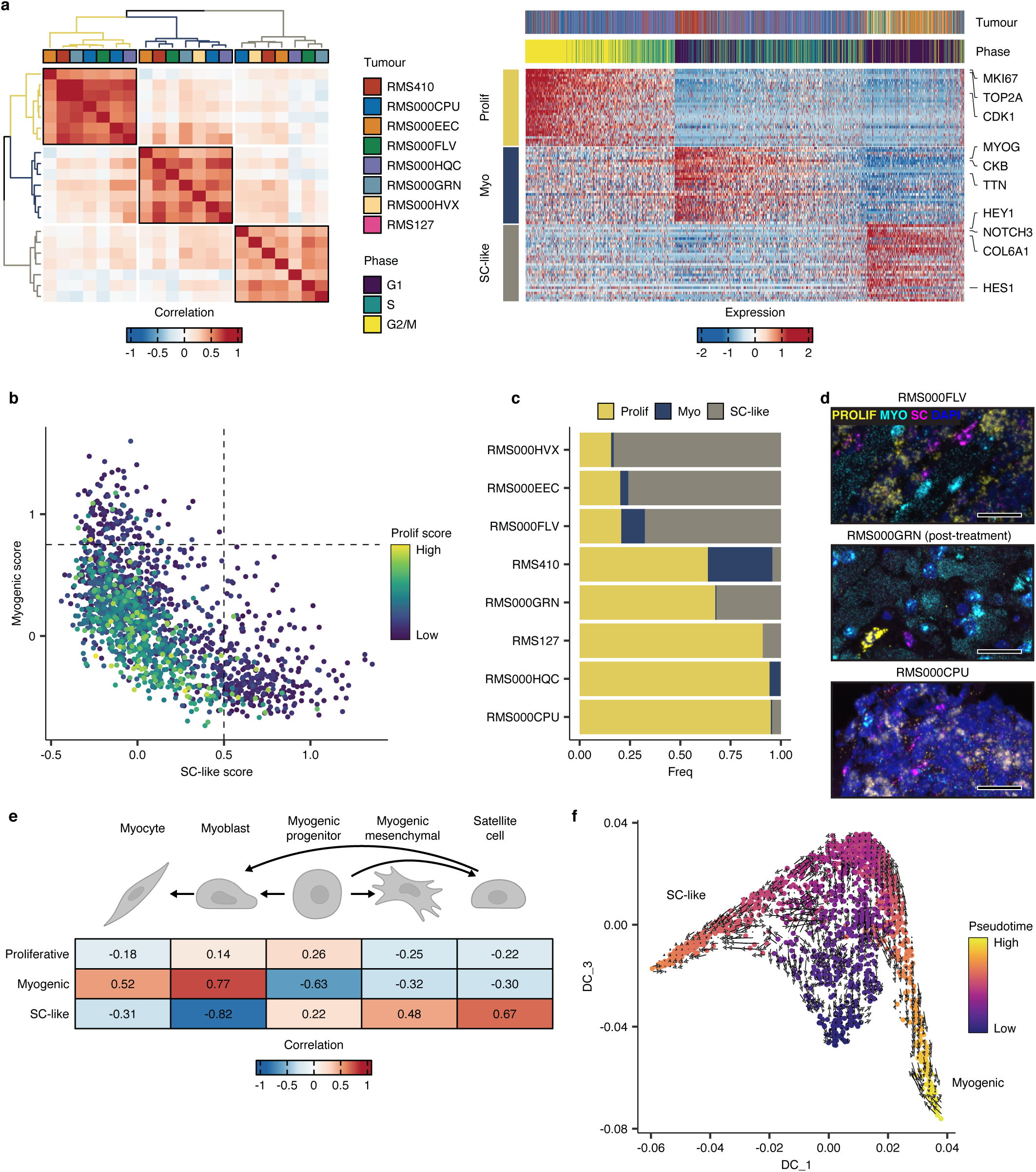
Cell states in FP RMS tumours mirror skeletal muscle myogenic differentiation. **a**, Left panel: Heatmap showing the pairwise Pearson correlations between all NMF-defined transcriptional programs in FP samples. The tumour sample from which each transcriptional program derived is shown in the colour bar. Meta-program clusters are delineated by black boxes and colouring of the dendrograms. Right panel: Scaled expression of the top 30 genes per meta-program across all FP cells. The corresponding tumour sample and inferred cell cycle phase of each cell are displayed in the top annotation bar. Representative genes from each meta-program are labelled. **b**, Scatterplot depicting per cell meta-program scores. Dotted lines correspond to the cut-offs used to define discrete cell states. **c**, Proportion of cells within each discrete state, per FP tumour. **d**, Representative RNA fluorescence *in-situ* hybridization (RNA-FISH) images depicting the expression of satellite cell-like (magenta, SC = *NOTCH3*), myogenic (cyan, MYO = *TTN*) and proliferative (yellow, PROLIF = *MKI67*) cell state marker genes in FP tissue samples. DAPI counterstaining shown in blue. Scale bars equivalent to 25*µ*m. **e**, Heatmap depicting the Pearson correlations between FP cell-state scores, and the logistic regression-defined similarity scores (logits) for each normal myogenic cell type. **f**, Diffusion maps projection of FP RMS single cells, coloured by pseudotime value, overlaid with the RNA velocity vector field. Myogenic differentiation schematic was created with BioRender.

As with the FN samples, there was a high degree of variation in discrete cell-state proportions between tumours, particularly among the proliferative and SC-like states (Fig. 4c). The expression of each meta-program, as well as the mutual exclusivity of the myogenic and SC-like programs was validated using RNA-FISH (Fig. 4d). Comparisons between meta-program scores and the logistic regression-defined cell similarity scores showed that the myogenic program correlated strongly with cell types along the terminal differentiation trajectory (myoblasts and myocytes) while the SC-like program was comparable to post-natal satellite-cells (Fig. 4e). Though the proliferative program score did not strongly correlate with any of the normal myogenic cell types, supporting the notion that this program was indicative only of cell cycle activity, most cells within the proliferative state most closely resembled myoblasts (Extended Data Fig. 6c). Trajectory inference indicated that cells scoring high for the myogenic or SC-like programs lay at opposite ends of the differentiation continuum, while the proliferative cells appeared as an undifferentiated intermediate state (Fig. 4f). In this case, however, the RNA velocity results did not definitively imply a strict directionality of the trajectory (Fig. 4e). Altogether, these data showed that the shared cell-state heterogeneity in FP RMS forms a differentiation trajectory reminiscent of that underlying skeletal muscle regeneration, where SC-like cells connect to cells resembling proliferative, undifferentiated myoblasts, which may give rise to (or derive from) cell bearing similarity to terminally differentiating myoblasts/myocytes.

### Malignant cell states are predictive of patient outcomes

Taken together, results from the analysis of NMF-defined transcriptional programs allowed us to propose a unified model of cell states and differentiation trajectories in FN and FP RMS tumours (Fig. 5a). In FN tumours, highly proliferative cells with characteristics of early myogenic progenitors (progenitor-like) seem to give rise to cells which resemble either of two more differentiated types: myogenic mesenchymal cells (mesenchymal) or terminally differentiating myoblasts/myocytes (myogenic). On the other hand, in FP tumours, highly proliferative cells resembling committed myoblasts (proliferative) are an intermediate between cells closely resembling differentiating myocytes (myogenic), or post-natal muscle resident satellite cells (SC-like). To investigate whether the differentiation state of RMS tumours affects their clinical behaviour, a published cohort of bulk tumour gene expression profiles^8^ was scored for each meta-program. Strikingly, FN RMS patients whose tumours had a high differentiation score (mesenchymal + myogenic) exhibited a significantly better OS probability than those with a low score (*p* = 0.00069, Fig. 5b-left panel). This result was particularly intriguing, as neither cell-state program was predictive of outcomes on its own (Extended Data Fig. 7a,b).Conversely, a high score for the undifferentiated progenitor-like program was indicative of significantly worse OS than FN tumours with a low score (*p* = 0.035, Fig. 5b-right panel). In FP RMS patients, high expression of the SC-like program was associated with extended OS (*p* = 0.017), while a high proliferative score was indicative of shorter OS (*p* = 0.029, Fig. 5c). Differential expression of the myogenic program in FP tumours was not predictive of patient survival (Extended Data Fig. 7c). These data show for the first time that, in both RMS subtypes, tumours with higher proportions of cells in more differentiated states exhibit better outcomes than those with high levels of proliferative, less well differentiated cells.

**Figure 5:**
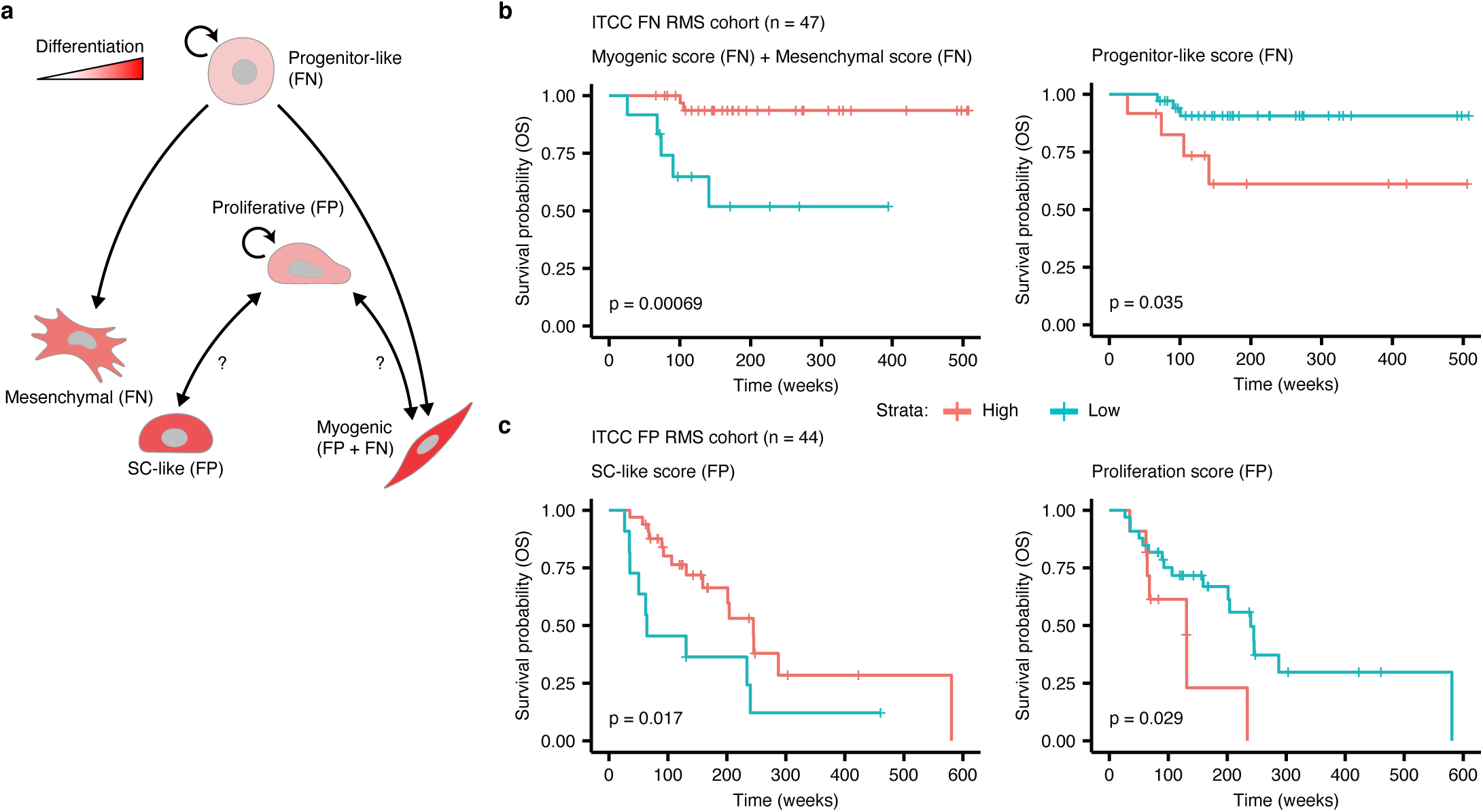
Malignant cell states are predictive of patient outcomes. **a**, Schematic representation of differentiation trajectories in RMS. Created with BioRender. **b**,**c**, Kaplan-Meier plots showing the overall survival probabilities of (**b**) FN or (**c**) FP patients divided into high (red strata) or low (blue strata) groups based on their cell state scores (stated in the title of each plot panels). Log rank test was used to calculate *p* values between high and low scoring groups.

## Discussion

We generated a single-cell transcriptomic atlas of RMS tumours, detailing cell states of both malignant cells and those constituting the TME. In our investigation of the TME, we found that among differentiated macrophages in both RMS subtypes, the immunosuppressive, pro-tumour M2-type was predominant. This result differs from the findings in two recent publications, which found a roughly balanced proportion of M1 and M2 macrophages in both RMS subtypes^31,32^. This deviation may be due to the fact that in both studies, single markers were used to delineate M1 and M2-type macrophages (CD68 and CD163, respectively), while the analysis presented here relied on multi-gene signatures which may be more robust in characterising cell states. We also described, for the first time, a putative interaction between FP tumour cells expressing NECTIN3 and the TIGIT receptor on Tregs and NK cells, which we validated using immunofluorescence staining of patient tissue samples. This interaction may result in the suppression of anti-tumour immune responses through several mechanisms, as has been described in other malignancies^33^. As such, targeting this interaction could represent an opportunity to sensitize FP RMS tumours to immune-mediated killing through blocking of the TIGIT receptor, an approach which is currently being clinically evaluated for other tumour types^33^. Overall, we observed a higher proportion of T/NK cells, relative to myeloid cells (∼1:1), than has been previously described in studies utilizing immunohistochemistry^31,32^ or scRNA-seq^28^. Additionally, beyond endothelial cells, we were unable to detect any other non-immune cell types in the TME, such as cancer associated fibroblasts (CAFs) or non-malignant skeletal muscle cells. We ascribe these inconsistencies to either biases introduced by the SORT-seq protocol, (multi-nucleated myotubes would be gated out during FACS sorting, for instance) or the freeze/thaw cycle tumour samples were subjected to. However, including a larger number of T and NK cells had the benefit of allowing us to resolve and characterize functional subtypes not previously identified in RMS tumours.

In analysing NMF-defined transcriptional meta-programs and the similarity of RMS single-cells to normal myogenic differentiation, we defined subtype-specific hierarchies of malignant cell states. While myogenic differentiation of RMS cells into rhabdomyoblasts has long been appreciated^3^, the presence of cells resembling myogenic mesenchymal cells or satellite cells has, to our knowledge, not been previously described. Our model of RMS differentiation trajectories (Fig. 5a) has several clinical and biological implications. First, the observation that high levels of cells in more differentiated states is associated with better patient outcomes suggests the use of “differentiation therapy”, where tumour cells are pharmacologically induced to undergo differentiation^34^, would be a useful treatment strategy for RMS. In support of this, several studies using pre-clinical models of RMS have demonstrated that inhibiting critical pathways or regulators of tumorigenesis, including MEK in mutant RAS-driven FN RMS^35^ and BAF complexes in FP tumours^36^, leads to the induction of terminal myogenic differentiation. This approach could be expanded upon in future studies through the systematic elucidation of key regulators of RMS cell states which could be targeted to induce differentiation. It was notable that in both RMS subtypes, high levels of cells in states with mesenchymal characteristics (FN mesenchymal and FP SC-like) were associated with better outcomes, as mesenchymal-like cell states have been associated with drug resistance and increased metastatic potential in tumours of epithelial origin^37^ as well as other sarcoma types^38^. On the other hand, the observation that high levels of proliferation are associated with worse outcomes supports the potential utility of compounds targeting key cell cycle regulators, including WEE1, PLK1 or CDK4/6 inhibitors, all of which are being investigated as therapeutic additions to RMS treatment regimens^39^. In addition to informing future treatment strategies, our results suggest that the differentiation state of RMS tumours could be a valuable metric for patient stratification, particularly in FN RMS. The translation of this finding could help advance a key goal of RMS clinical research: the de-intensification of treatment, where possible, to reduce toxicity and treatment-induced late effects^40^. However, these results first need to be validated using independent patient cohorts, an effort which is complicated by the overall lack of publicly available data sets combining gene expression of RMS tumours with clinical follow-up information. Finally, while RNA velocity analysis suggested that the myogenic, mesenchymal, and SC-like states derive from the more proliferative cell states, it does not definitively rule out the possibility of de-differentiation from more to less-well differentiated cell states. It will be important, therefore, in future studies to examine the dynamic relationships between cell states using, for instance, phylogenetic analyses^41^ or functional assays in pre-clinical RMS models.

Notably, our proposed model of differentiation trajectories in RMS differs from that recently put forth by Patel *et al* ^28^. In their analysis, a single trajectory explains both FN and FP RMS, whereby cells resembling paraxial mesoderm (*MEOX2*+, found mainly in FN tumours) connect to highly proliferative myoblast-like cells which in turn may give rise to (or derive from) a more differentiated myocyte-like state. This difference in interpretation could stem from the fact that Patel *et al*, jointly analysed FN and FP samples, which may have obscured subtype-specific differences. For example, while highly proliferative myoblast-like cells were observed in our FP samples, the highly proliferative state in FN tumours more closely resembled early myogenic progenitors. However, as we have shown in Extended Data Figs. 5b and 6b, the data from Patel *et al* was consistent with our subtype-specific models of RMS differentiation trajectories.

In our comprehensive analysis of single-cell transcriptomes from paediatric RMS, we characterized the immune component of the TME and defined cell-states mirroring normal myogenic differentiation trajectories. Based on these findings, we propose that targeting immune checkpoint molecules is a promising therapeutic approach for RMS that merits further investigation. Furthermore, the validation and clinical implementation of differentiation state as a prognostic indicator should be a priority, given its potential to improve patient risk stratification.

## Methods

### Tumour sample acquisition

Tumour samples of RMS were obtained via an established sample acquisition route as part of the biobank initiative of the Princess Máxima Center for Pediatric Oncology, Utrecht, Netherlands (remaining tumour samples). Ethics approval was granted for the biobanking initiative by the Medical Research Ethics Committee (METC) of the University Medical Center Utrecht, and the Maxima biobank committee granted approval for the present project. All patients and/or their legal representatives signed informed consent to have tumour samples taken for biobank usage. Experiments conformed to the principles set out in the WMA Declaration of Helsinki and the Department of Health and Human Services Belmont Report.

### Sample processing and single-cell RNA-sequencing

Viably frozen tumour samples were rapidly thawed in a water bath, minced using a scalpel and then transferred to a tube containing 4.5 ml of BM1* medium (Advanced DMEM/F12 [Gibco, cat no. 12634010] supplemented with 1% Glutamax [Gibco, cat no. 35050061], 1% Penicillin/Streptomycin [Gibco, cat no. 15140122], 2% B27 minus vitamin A [Gibco, cat no. 12587010], 1% N2 [Gibco, cat no. 17502048], 0.25% N-acetylcysteine [500 mM, Sigma, cat no. A9165], 1% MEM non-essential amino acids [Gibco, cat no. 11140035], 1% sodium pyruvate [100 mM, Gibco, cat no. 11360070], 0.01% heparin [5,000 U/ml, Sigma, cat no. H3149-10KU], 1% hEGF [2 *µ*g/ml, Peprotech, cat no. AF-100-15], 0.1% hFGF-basic [40 *µ*g/ml, Peprotech, cat no. 100-18B], 0.02% hIGF1 [100 *µ*g/ml, Peprotech, cat no. 100-11], 0.01% Rho kinase inhibitor [Y-27632, 100 mM, AbMole Bioscience, cat no. M1817] and 0.1% A83-01 [5 mM, Tocris Bioscience, cat no. 2939]). To this, 0.5 ml of Collagenase D (Roche, #11088866001, 1:10 dilution) and DNAseI (Stemcell #07900, stock diluted 1:40 in PBS, further 1:100 diluted in the BM1* mixture) were added, and samples were allowed to dissociate in a shaker set to 250 rpm for 30 minutes at 37°C. Following digestion, samples were passed through a 70 *µ*m strainer which was subsequently flushed with an additional 5ml of BM1* (supplemented with DNAseI) to increase the yield. Samples were then washed twice with 5ml of washing medium (Advanced DMEM/F12 supplemented with 1% Glutamax, 1% Penicilin/Streptomycin and 1% HEPES [1M, Gibco, cat no. 15630049]), centrifuging at 300g for 5 minutes (at 4°C) in between steps. After the final washing step samples were resuspended in BM1* (supplemented with DNAseI) to a final concentration of < 1 × 10^6^ cells per ml. Prior to sorting, 4′,6-diamidino-2-phenylindole (DAPI, Sigma-Aldrich, #D9542) and DRAQ5 (Thermo Fisher, #65-0880-92) were added to single-cell suspensions up to final concentrations of 1*µ*M and 5*µ*M, respectively. Viable single-cells (DAPI-, DRAQ+) were then sorted into 384-well plates containing 10*µ*l of mineral oil (Sigma, #M5310) and 50nl of barcoded RT primers using a SONY SH800S Cell Sorter. Libraries were prepared according to the SORT-seq^15^ protocol and sequenced on an Illumina NextSeq500 (paired-end, 75bp read chemistry) by Single Cell Discoveries B.V..

### Immunohistochemistry and H&E staining

Immunohistochemistry (IHC) and haematoxylin and eosin (H&E) staining experiments were performed on 4*µ*m thick formalin fixed and paraffin embedded (FFPE) tissue sections using a Ventana automated tissue staining system (BenchMark Ultra, Roche). For IHC, the antibodies used were anti-CD3 clone LN10 (Leica, PA0533), anti-CD8 4B11 (Leica, PA0183) and anti-CD68 514H12 (Leica, PA0273).

### Immunofluorescence microscopy

Mounted tumour sections (5 *µ*m thick FFPE) were baked at 60°C for 1 hour and then deparaffinized and rehydrated using sequential washes of Xylene (2 × 100%), Ethanol (2 × 100%, 2 × 95%, 1 × 75%, 1 × 50% and 1 × 25%) and demineralized H2O (2 × 1 minute, 1 × 5 minutes). Antigen retrieval was then performed by boiling slides in Tris-EDTA (pH 9) for 20 minutes in a benchtop autoclave. Slides were then washed 3×5 minutes in PBST (PBS + 0.1% Tween 20) and incubated with blocking solution (PBST + 1% BSA) for 1 hour at room temperature. After blocking slides were incubated with primary antibody diluted in blocking solution overnight at 4°C. The following day, slides were washed 3 × 5 minutes with PBS and then incubated with secondary antibody, diluted in PBST, in the dark for 1 hour at room temperature. Slides were washed an additional 3 × 5 minutes with PBS before adding mounting medium containing DAPI counterstain (Vector labs, H-1200) and applying glass coverslips. Images were acquired on a Leica SP8 confocal microscope (40x/1.3NA oil immersion objective), and maximum projections of Z-stacks were obtained using the FIJI software (v2.0.0-rc-69/1.52i) ^42^. Primary antibodies used: anti-TIGIT (Cell Signaling, #99567, 1:500 dilution) and anti-NECTIN3 (R&D systems, AF3064, 1:200 dilution of a 0.2*µ*g/*µ*l solution in PBS). Secondary antibodies used: Donkey anti-Goat Alexa 647 (Abcam, ab150131, diluted 1:1000) and Donkey anti-Rabbit Alexa 568 (Abcam, ab175470, diluted 1:1000).

### RNA Fluorescence *in-situ* hybridization (RNAscope)

RNA-FISH experiments were performed on 5 *µ*m FFPE tissue sections using the RNAscope™ Multiplex Fluorescent v2 kit (ACD bio), according to the manufacturer’s instructions. The following probes were used for hybridization: Hs-MKI67-C3 (591771-C3), Hs-TTN (550361), Hs-NOTCH3-C2 (558991-C2), Hs-FGFR4-no-XMm-C2 (443431-C2) and Hs-TGFBI (478491). Additionally, the following fluorescent dyes were used for detection (diluted 1:1500): Opal 520 (FP1487001KT), Opal 570 (FP1488001KT) and Opal 690 (FP1497001KT). Images were acquired on a Leica SP8 confocal microscope (40x/1.3NA oil immersion objective), and maximum projections of Z-stacks were obtained using the FIJI software (v2.0.0-rc-69/1.52i).

### Data processing and quality control

Sequencing reads were demultiplexed, mapped to the GRCh38v2020-A genome, available from 10X genomics (https://support.10xgenomics.com/single-cell-gene-expression/software/release-notes/build), and transcript counts were generated using the zUMIs pipeline (v5.6)^43^. Using the Seurat toolkit (v4.1.0)^44^, count tables (per plate) were then loaded in R (v4.1.0), merged and metadata fields were compiled. Single cells excluded if they had < 500 expressed genes, < 800 or > 50,000 unique transcripts, a percentage of mitochondrial transcripts > 50%, > 1% haemoglobin gene transcripts or a ratio of intergenic to genic transcripts > 2. The data were then log normalized to 10,000 transcripts, scaled and centered. The top 2,000 most variably expressed genes were defined with the *FindVariableFeatures* function in Seurat (default parameters), and their expression was used as input for principal component analysis (PCA). Finally, the first 50 principal components were used to project single-cell transcriptomes in 2-dimensional space using uniform manifold approximation and projection (UMAP). The cell cycle phase of each single-cell was inferred using the *CellCycleScoring* function implemented in Seurat, using the built-in gene lists.

### Module scoring

Module scores were calculated as previously described in ref. ^20^ and implemented in the Seurat function *AddModuleScore*, taking into account 25 expression bins and 100 control genes per query gene.

### Cell type classification

To discriminate between malignant and healthy cells, we first used the SingleR R package (v1.6.1)^16^ to annotate single cells based on their similarity to reference bulk transcriptomes of healthy cells (Human Primary Cell Atlas data^45^) and RMS tumours (EGAD00001008467). We then used the InferCNV R package (v1.8.0)^46^ to define and cluster single-cell copy number variant profiles per tumour sample (default parameters, average expression threshold of 0.3 and standard deviation filter of 2). A SORT-seq dataset of cord blood mononuclear cells (CBMC’s) and other normal cell types was used as reference. CNV profiles derived from bulk DNA sequencing were plotted for comparison (see Extended Data Fig. 2), and single-cell clusters containing CNVs were manually selected and annotated “malignant”. Cells were called malignant when they were classified as such using both approaches and excluded cells which were divergently classified (labelled ambiguous). The broad cell-type of non-malignant cells was inferred from the hierarchical clustering of the similarity scores.

### Analysis of the immune microenvironment

To reach sharper biological distinctions between immune cell subsets, SCTransform^47^ normalization was performed on the full dataset to normalize and scale the data for unbiased clustering. To further improve detailed immune cell sub-clustering, sample-specific gene expression was removed to reduce technical effects and enhance biological variation. Sample-specific genes were identified by differential gene expression analyses among tumour cells and immune cells separately and comparing the individual samples. Genes that were differentially expressed in both the tumour cells and immune cells of a specific samples were considered sample-specific noise and were removed from the variable gene list. To avoid clustering of cells based on specific cell processes, genes associated with sex (*XIST, TSIX*, and Y chromosome-specific genes), cell cycle phase, dissociation stress (heat shock proteins; GO:0006986), and activity (ribosomal protein genes; GO:0022626), were also removed from the variable gene list as described previously^48^.

Healthy clusters were subset and clustered using 40 principal components and a resolution of 0.3 (Louvain algorithm) was used to define clusters of the main cell types. For in-depth analysis of the T and NK cells, the respective clusters were subset, and UMAP was re-run using 40 PCs and a resolution of 0.5 was used to define subclusters. For in-depth analysis of the myeloid compartment, SCTransfrom normalization was re-run, sample-specific and cell process-specific genes were removed from the variable gene list and 6 PCs and a resolution of 0.3 was used to define subclusters.

### Immune cell type identification

Cluster annotation was performed using SingleR, using the Human Primary Cell Atlas reference dataset to annotate main cell types, and additionally using the Novershtern Hematopoietic Data^49^ and Monaco Immune Data^50^ reference datasets to annotate the immune cell (sub)clusters. Cell annotations were further refined by consulting cluster-specific (up-regulated) differentially expressed marker genes using Seurat’s *FindAllMarkers* function. The outputted genes were compared to known cell-type specific marker genes from previous studies^51–54^.

### Gene set enrichment analysis (GSEA)

For GSEA, differential expression analysis between two groups was performed using the *FindMarkers* Seurat function, using the following adjusted parameters: logfc.threshold=0, min.pct = 0, min.cells.feature = 0, min.cells.group = 0. Genes were pre-ranked by their Fold Change and GSEA was performed using the R package fgsea (version 1.20.0). Gene sets with an FDR <0 .25 were considered significantly enriched. Gene sets were obtained from MSigDB version 7.2.

### Ligand-receptor interaction analysis

The CellChat^24^ algorithm was applied to perform an unbiased ligand-receptor interaction analysis, using the curated ligand-receptor database of CellPhoneDB (RRID: SCR_017054)^55^.

### Logistic regression analysis

Determination of the similarity between RMS single-cells and normal myogenic cells types (given as a probability value) was estimated as previously described in ref. ^26^. We obtained the data from ref. ^27^ from the gene expression omnibus (GSE147457) and trained logistic regression models using the main myogenic cell type labels. Correlations between meta-program scores and normal myogenic cell types used the logit-transformed probability values.

### Non-negative matrix factorization

Non-negative matrix factorization (NMF) was carried out using the NMF R package (v0.23)^56^. For each RMS subtype (FN or FP), a list of shared variable features (*n* = 2000) was compiled using the *SelectIntegrationFeatures* function in Seurat. The expression of these genes was then scaled, per tumour, and used as input to determine the appropriate NMF rank, by running 50 iterations (Brunet algorithm) for ranks between 2 and 10 (default settings). The optimal rank was determined, per tumour, by manually assessing in cophenetic coefficients, dispersion values and silhouette scores between rank values. We then re-performed NMF at 250 iterations using the optimal rank value. Per subtype, pairwise Pearson correlation coefficients were calculated between NMF-defined transcriptional programs (across all tumours) and hierarchical clustering was used to determine groups. Highly correlated groups of programs were merged into meta-programs by averaging gene weights. Cell-state scores were by using the top 30 weighted genes per meta-program to calculate module scores. Discrete cell-states were determined through manual inspection of the distribution of cell-state module scores.

### Gene list enrichment analysis

Functional enrichment of gene lists was performed by the enrichR R package (v3.0)^57^ (default settings) using the Reactome 2016 database.

### Comparison with data from *Patel et. Al*

Single-nucleus RNA-seq data from the manuscript of Patel *et. al*, was downloaded from the Single-Cell Pediatric Cancer Atlas Portal (https://scpca.alexslemonade.org/) and loaded into Seurat. We inferred malignant cells using SingleR, as described for the data presented in this paper and applied an additional cutoff of > 800 unique transcripts for a cell to be considered valid. Data were then split by molecular subtype and module scores were calculated, as described above, using the top 30 genes per meta-program.

### Differentiation trajectory modelling

Modelling of differentiation trajectories was done, per subtype, by projecting cells in DiffusionMaps space using expression of the top 30 meta-program-specific genes (destiny R package v3.1.1)^58^. The top 3 diffusion components were then used as input for trajectory modelling and cell lineage inference using Slingshot (v2.0.0)^59^. RNA velocity analysis was performed using the scVelo python package (v0.2.2, python v3.7)^60^. Briefly, input data per subtype, was filtered to include only genes with 20 shared (spliced and un-spliced) counts and log normalized. First and second order moments were calculated per cell using expression of the top 30 meta-program-specific genes and 30 nearest neighbours. RNA velocity was then estimated using the stochastic model and vectors were overlaid on the DiffusionMaps projections.

### Survival analysis

Microarray gene expression profiles and the accompanying clinical follow-up information for the ITCC RMS cohort^8^ was downloaded from the R2 genomics platform (R2: Genomics Analysis and Visualization Platform (http://r2.amc.nl)). Samples which did not exhibit either of the two main RMS histological classifications (alveolar or embryonal) were excluded. The data were divided based on fusion transcript status and Z-scores were calculated per gene. To generate meta-program scores, the average Z-score of the top 30 genes per meta-program (in the appropriate dataset) was calculated per tumour. Based on the distribution of scores, the “high” scoring groups (and vice versa) were defined as either the top 25% or 75% of tumours. Survival models were generated using the survival R package (v3.2-11) and *p* values were calculated using a Log Rank test.

## Data availability

Raw single-cell RNA sequencing data have been deposited in the European genome-phenome archive. Accession numbers are pending XXX.

## Code availability

All data analysis code will be made available without restriction upon request.

## Acknowledgements

We thank the patients and their families who made this research possible by consenting to this study. We also thank the many clinicians in our institute with whom we work closely, as well as the Princess Máxima Center’s flow cytometry, single-cell genomics and imaging core facilities. This research was supported by the Foundation Children Cancer-free (Kika) and a European Research Council (ERC) starting grant #850571 (J.Dr.). M.M. received financial support from the Deutsche Forschungsgemeinschaft (#408083583).

## Contributions

J.De., M.M., L.V., F.H., J. Dr. and T.M. conceived and designed the experiments and analyses. J.De. and L.V. analysed the data with contributions from J.Dr., T.S. and T.M. J.De., M.M. and M.G.K. processed tumour samples and performed FACS sorting for SORT-seq with contributions from T.M. L.H-J. provided detailed technical advice and acquired samples for the imaging experiments. J.De. and M.B performed the imaging experiments. J.De. wrote the manuscript with contributions from L.V., J.Dr., T.M., J.M. and F.H. J.Dr., T.M. and F.H. supervised the work. All authors read and approved the manuscript.

## Extended data figure legends

**Extended data figure 1.**
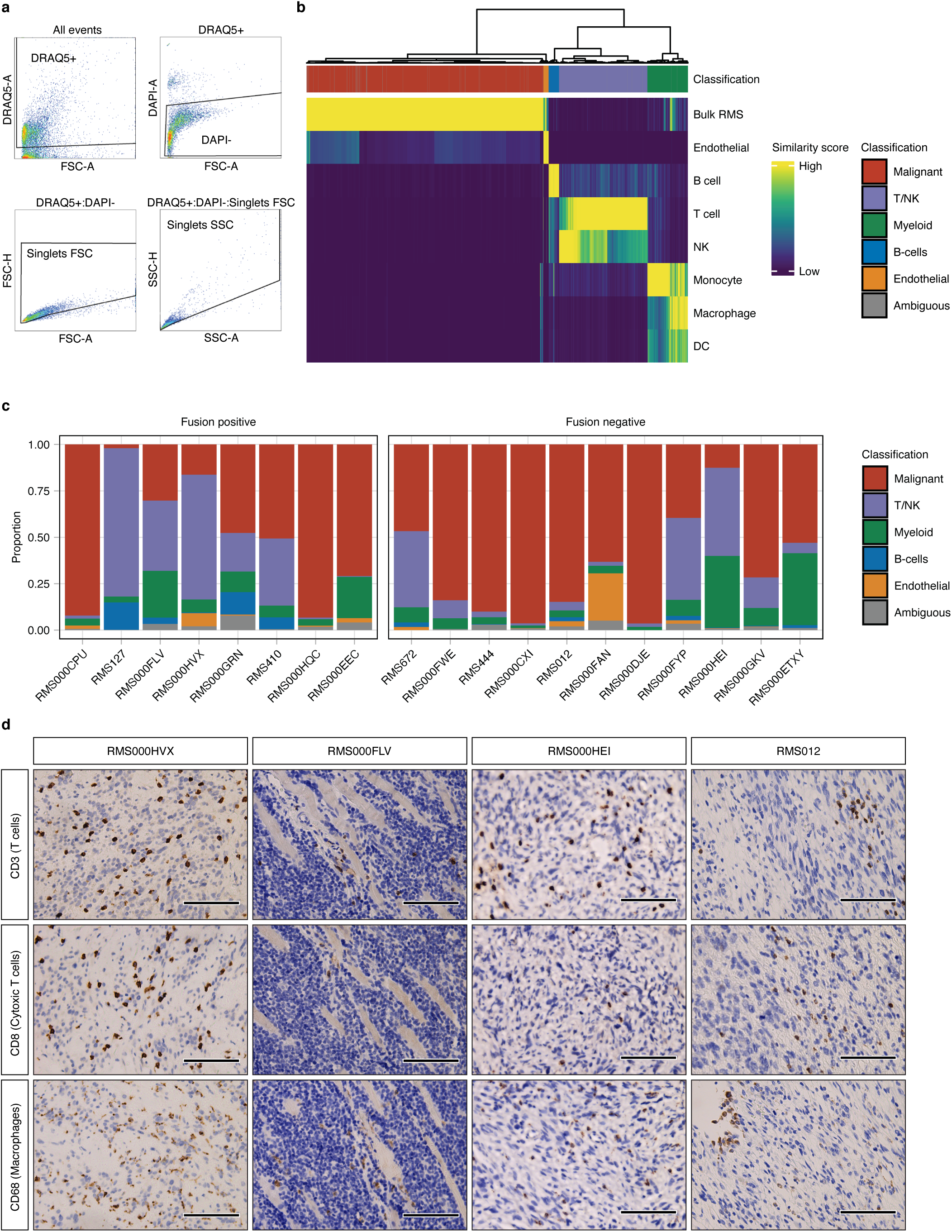
**a**, Representative scatter plots showing the gating strategy used for SORT-seq FACS. Titles indicate the events shown in each plot (FSC = forward scatter, SSC = side scatter, -A = signal area and -H = signal height). **b**, Heatmap showing the clustered similarity scores (per cell) to each reference cell type (y-axis labels) as determined by *SingleR*. Cell classifications are shown in the top annotation bar (as in Fig. 1). **c**, Bar graph depicting the distribution of annotated cell types per tumour. **d**, Representative microscopy images showing IHC staining of three immune population markers (y-axis) across tissue samples from four tumours (x-axis). Positive staining indicated by brown colouring, haematoxylin counterstained nuclei in blue. Scale bars equivalent to 100*µ*m.

**Extended data figure 2.**
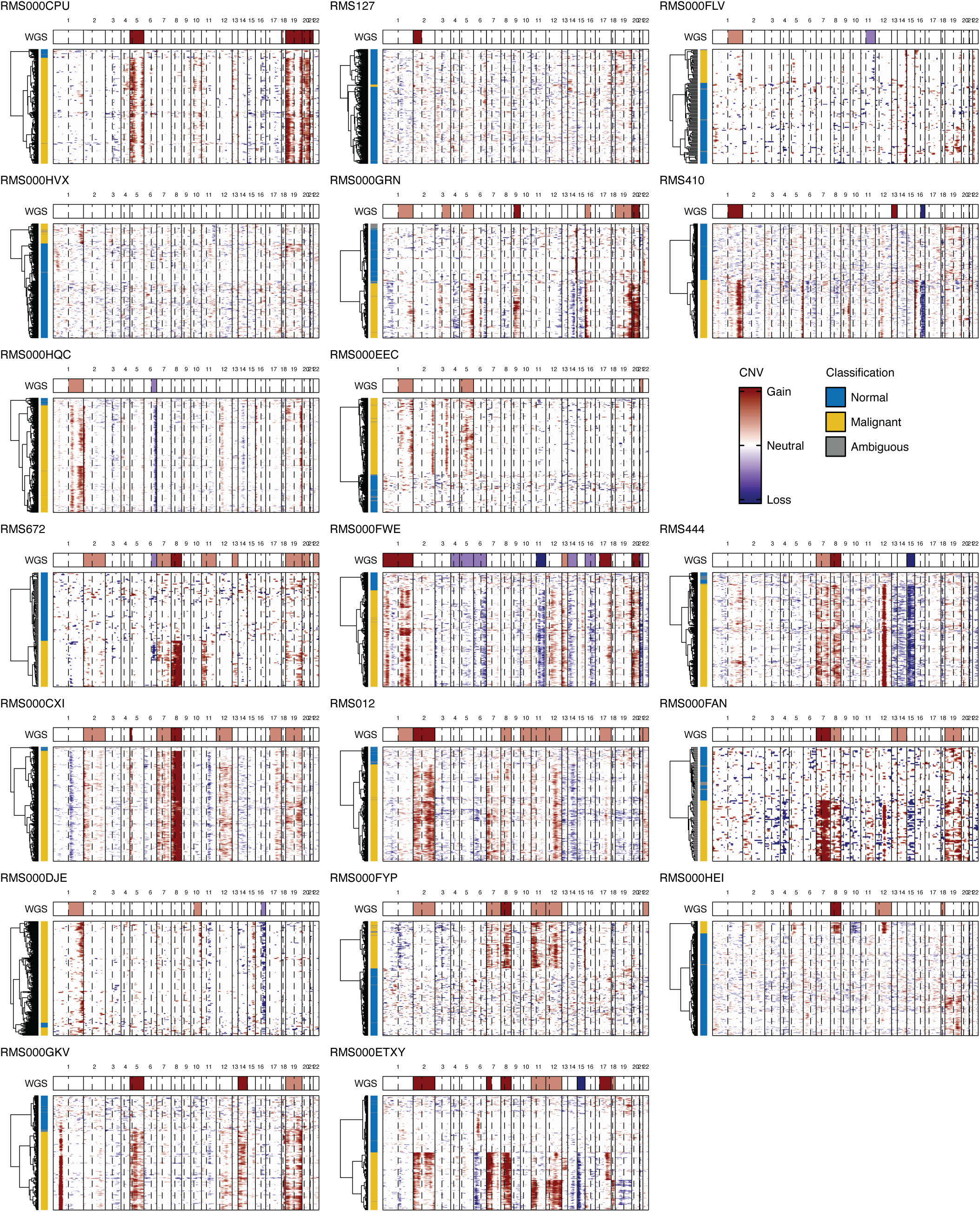
Heatmaps showing the clustered single-cell inferred CNV profiles per tumour. Solid vertical lines denote chromosome boundaries and dotted vertical lines represent the locations of centromeres. Cell classifications are shown on the left annotation bars. Top annotation bars show CNVs (summarized per chromosome arm) defined by DNA-sequencing of bulk tumour samples, for comparison.

**Extended data figure 3.**
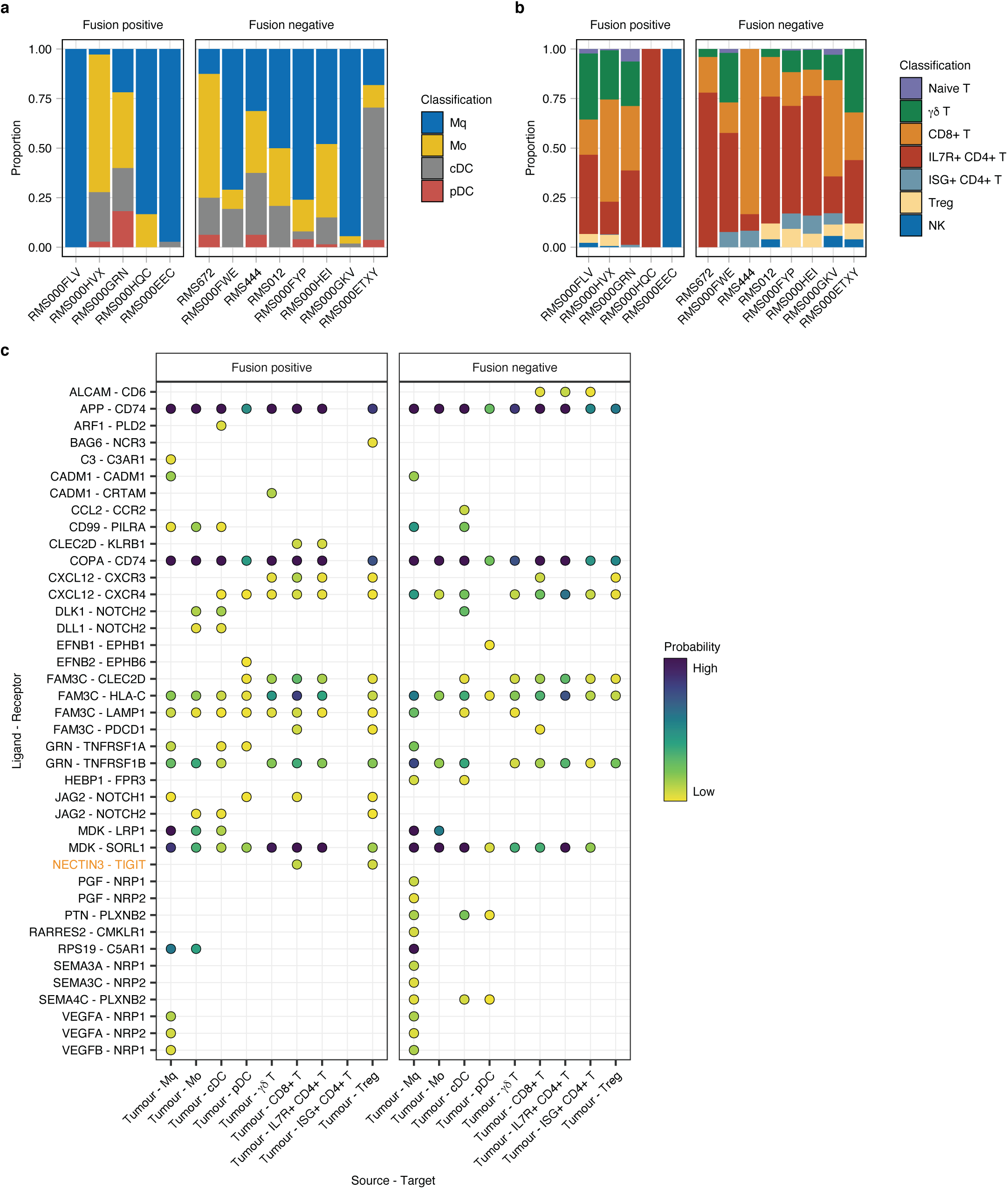
Bar plots showing the proportion of cell types within the (**a**) myeloid or (**b**) T/NK compartments per tumour. **c**, Dot plot summarizing the results of ligand-receptor interaction analysis, split by molecular subtype. Dots indicate an inferred ligand-receptor interaction (y-axis) between a source-target cell type pair (x-axis). Dots are coloured by the interaction probability, as determined by *CellChat*. The NECTIN3-TIGIT interaction is highlighted in orange on the y-axis.

**Extended data figure 4.**
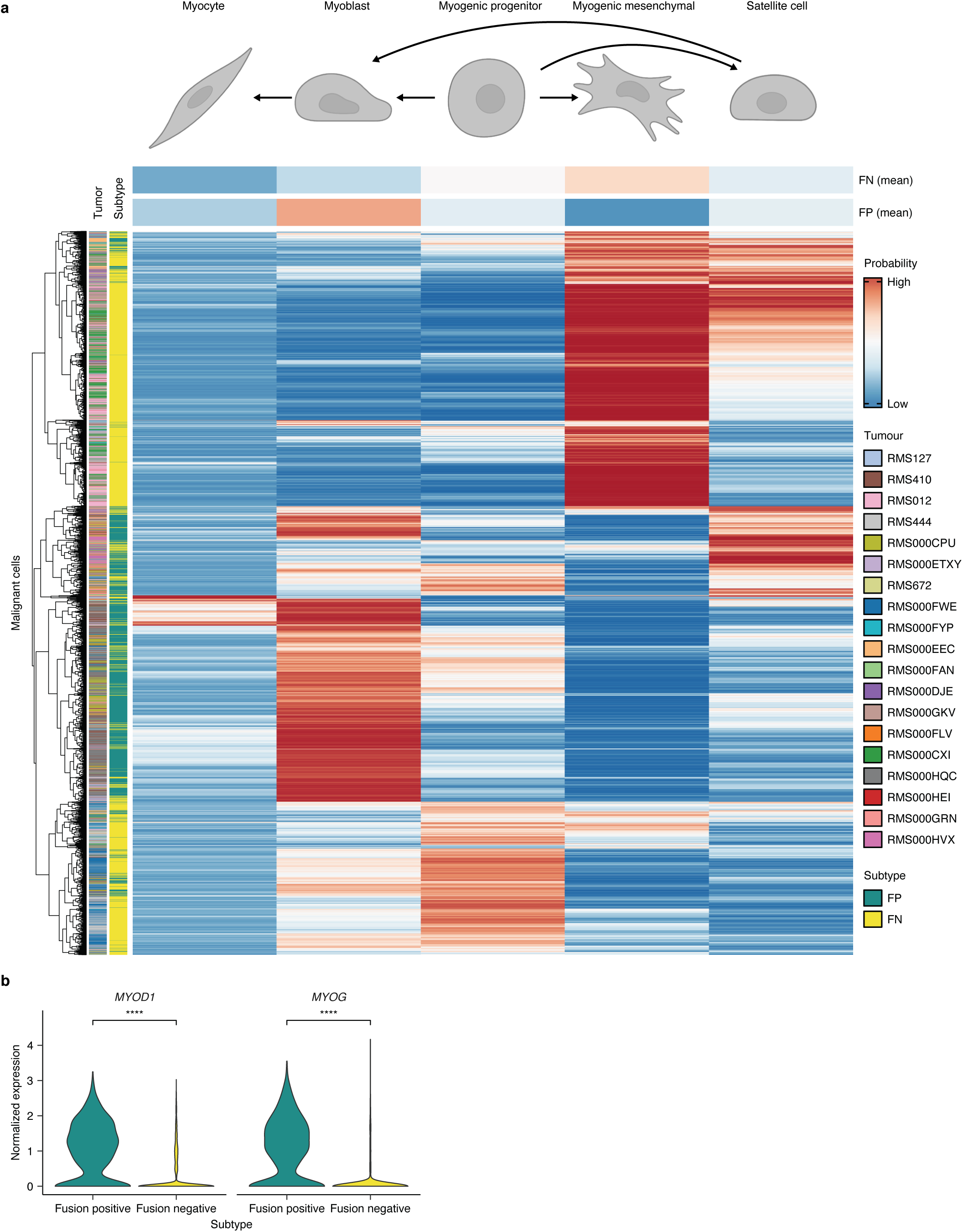
**a**, Heatmap showing the comparison between RMS single cells (rows) and normal myogenic cell types (columns). Colour values represent the predicted similarity (probability) as determined by logistic regression analysis. Annotation tracks (left) indicate the tumour and molecular subtype of each cell. The top two columns show the predicted similarity aggregated per molecular subtype. Myogenic differentiation schematic was created with BioRender. **b**, Violin plots depicting the normalized expression of *MYOD1* (left panel) and *MYOG* (right panel) in malignant cells from each RMS subtype. **** indicates differential expression (*p* < 0.00001, Wilcoxon Rank Sum test).

**Extended data figure 5.**
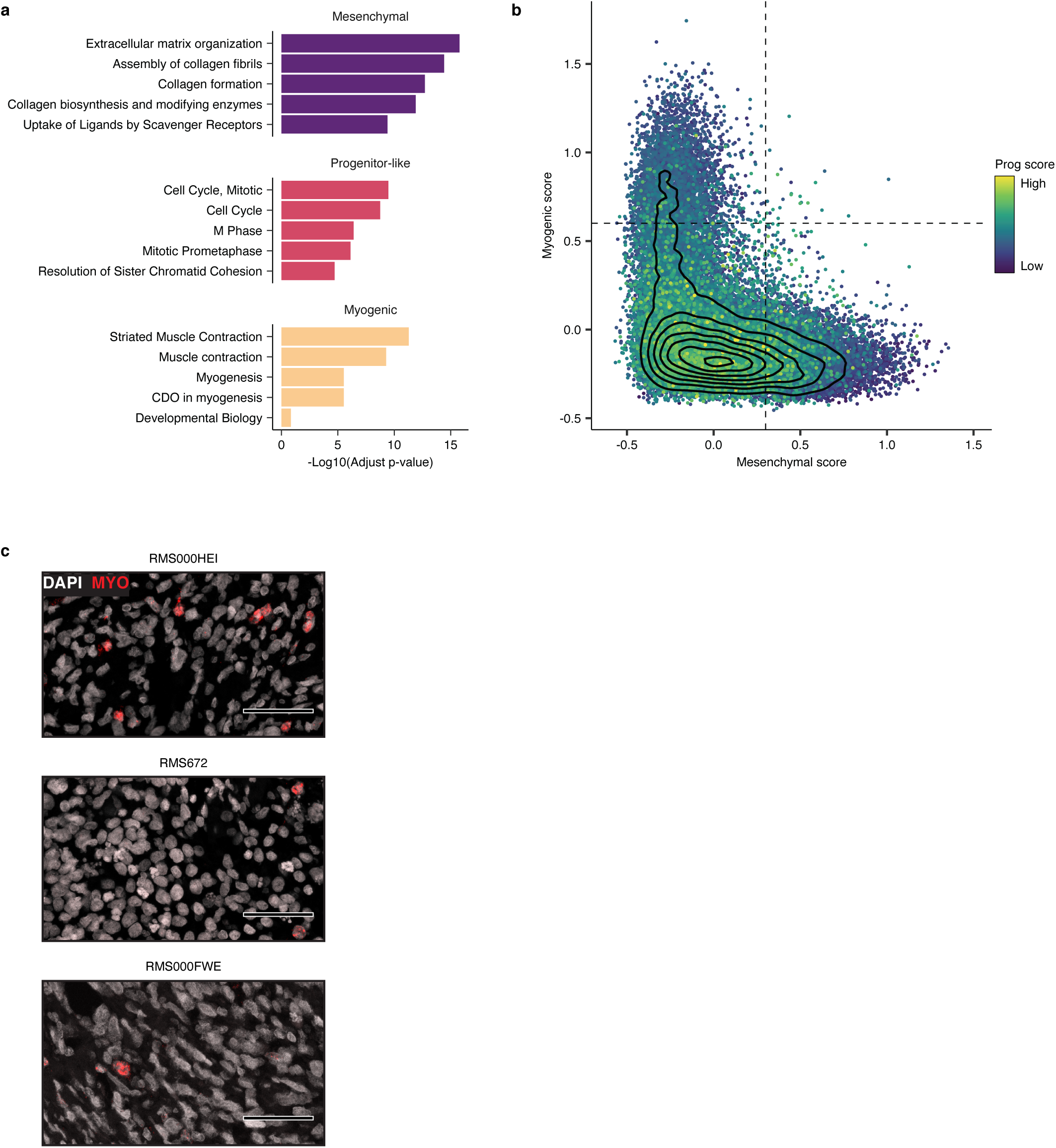
**a**, Reactome pathway enrichment of the top 30 genes per FN metaprogram. The top 5 terms per metaprogram are labelled on the y-axis and the -Log10-transformed adjusted *p*-values (Fisher’s exact test, B-H adjusted), are depicted by horizontal bars (coloured per meta-program). **b**, Scatter plot depicting the mesenchymal (x-axis), myogenic (y-axis) and progenitor-like (colour) meta-program scores calculated in the dataset from Patel *et. al* (malignant FN cells)^28^. Vertical and horizontal lines depict the discrete cell-state cut-offs used in Fig. 3b. Density contours are overlaid in black. **c**, Representative RNA-FISH images depicting the expression a myogenic cell-state marker gene in red (MYO = *TTN*) in FN tissue samples. DAPI counterstaining shown in grey. Scale bars equivalent to 50*µ*m.

**Extended data figure 6.**
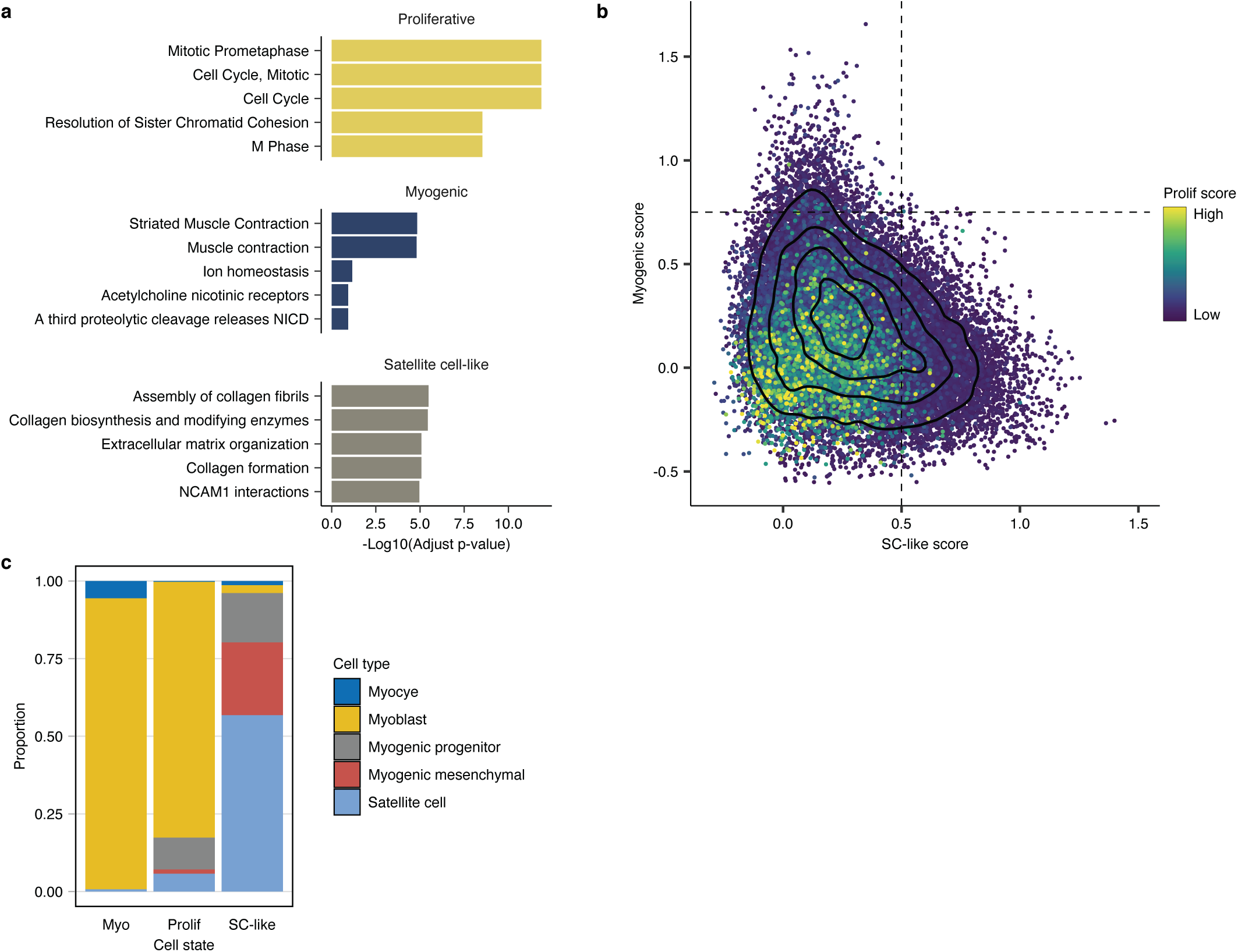
**a**, Reactome pathway enrichment of the top 30 genes per FP metaprogram. The top 5 terms per metaprogram are labelled on the y-axis and the -Log10-transformed adjusted *p*-values (Fisher’s exact test, B-H adjusted) are depicted by horizontal bars (coloured per meta-program). **b**, Scatter plot depicting the satellite cell-like (x-axis), myogenic (y-axis) and proliferative (colour) meta-program scores calculated in the data from Patel *et. al* (malignant FP cells)^28^. Vertical and horizontal lines depict the discrete cell-state cut-offs used in Fig. 4b. Density contours are overlaid in black. **c**, Bar graph showing the distribution of normal myogenic cell-type classifications (max probability score) for cells in each FP cell state (x-axis).

**Extended data figure 7.**
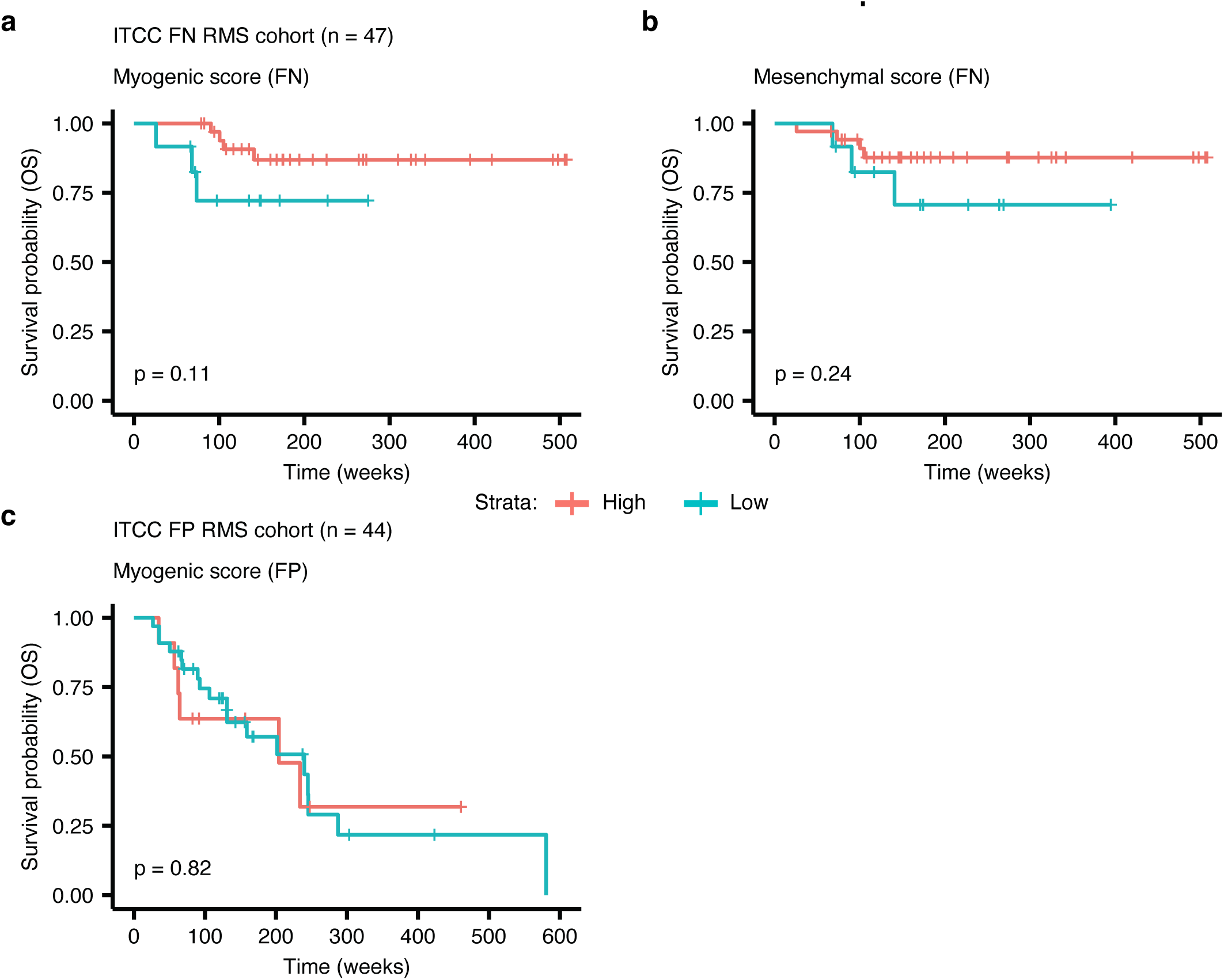
Kaplan-Meier plots showing the overall survival probabilities of (**a**,**b**) FN or (**c**) FP patients divided into high (red strata) or low (blue strata) groups based on their cell state scores (stated in the title of each plot panels). Log rank test was used to calculate *p* values between high and low scoring groups.

## Supplemental table legends

Supplemental Table S1: Detailed overview of RMS patient and sample characteristics

Supplemental Table S2: FN meta-program gene weights and top 30 genes

Supplemental Table S3: FN meta-program gene weights and top 30 genes

